# Taxonomic identification of airborne pollen from complex environmental samples by DNA metabarcoding: a methodological study for optimizing protocols

**DOI:** 10.1101/099481

**Authors:** Kleopatra Leontidou, Cristiano Vernesi, Johannes De Groeve, Fabiana Cristofolini, Despoina Vokou, Antonella Cristofori

## Abstract

Metabarcoding is a promising DNA-based method for identifying airborne pollen from environmental samples with advantages over microscopic methods. This method requires several preparatory steps of the samples, with the extraction protocol being of fundamental importance to obtain an optimal DNA yield. Currently, there is no consensus in sample preparation and DNA extraction, especially for gravimetric pollen samplers. Therefore, the aim of this study was to develop protocols to process environmental samples for pollen DNA extraction and further metabarcoding analysis, and to assess the efficacy of these protocols for the taxonomic assignment of airborne pollen, collected by gravimetric (Tauber trap) and volumetric samplers (Burkard spore trap). Protocols were tested across an increasing complexity of samples, from single-species pure pollen to environmental samples. A short fragment (about 150 base pair) of chloroplast DNA was amplified by universal primers for plants (trnL). After PCR amplification, amplicons were Sanger-sequenced and taxonomic assignment was accomplished by comparison to a custom-made reference database including chloroplast DNA sequences of 46 plant families, including most of the anemophilous taxa occurring in the study area (Trentino, Italy, Eastern Italian Alps). Using as a benchmark the classical morphological pollen analysis, it emerged that DNA metabarcoding is applicable efficiently across a complexity of samples, provided that sample preparation, DNA extraction and amplification protocols are specifically optimized.

## 1 Introduction

Pollen data retrieved by aerobiological surveys can provide information on several aspects, such as revealing flowering patterns of allergenic taxa, reconstructing the vegetation spectrum in a catchment area and studying the relationship of airborne pollen with vegetation and climate. Such information is important for both public health, since the revealed patterns can be related to seasonal allergies or for detecting impacts of climate change on plants (Damialis et al. 2007).

Airborne pollen surveys are conducted by means of volumetric or gravimetric methods, which collect pollen (and fungal spores) on different acceptors. In the case of the Hirst-type volumetric sampler, the most commonly used in aerobiological monitoring, pumped air directs pollen grains towards an adhesive surface, where they are trapped by impaction. Conversely, in the case of gravimetric sampling, pollen grains settle by gravity on a surface or into a sampling solution (e.g. Tauber traps) (Levetin 2004). In the current aerobiological studies, the sampling medium is then analysed microscopically in order to identify pollen taxa and define a pollen spectrum for the catchment area.

Few pollen grains can be identified at the species level by microscopic analysis because common morphological features are sometimes shared within genera and families and even between families. This is the case of Cupressaceae representatives sharing the same pollen morphological features with those of the Taxaceae family, leading to a microscopic classification of Cupressaceae-Taxaceae type. Microscopic analysis is also problematic in large-scale studies, since it requires considerable time and expertise for pollen identification and enumeration (Longhi et al. 2009).

Alternative methods for pollen identification have been investigated (e.g. spectroscope, optical-based detectors, etc.; Dell’Anna et al. 2010, Oteros et al. 2015). DNA-based methods appear particularly promising, especially thanks to the recent advent of Next Generation Sequencing (NGS). DNA metabarcoding of environmental samples has been already suggested as an alternative or complementary method to traditional approaches in fields related to pollen analysis, such as palaeopalynology, melissopalynology, diet assessment and biodiversity estimation (Parducci et al. 2013, De Barba et al. 2014). This approach identifies multiple species from a single complex environmental sample, targeting a specific part of DNA sequences (barcodes), which is shared across taxa, showing at the same time enough variation to allow taxonomic identification. It is worth noting that any metabarcoding approach has to be based on a comprehensive reference database, made after verified barcode sequences from the main taxa of the research area (Parducci et al. 2013).

DNA analysis on environmental air samples is particularly difficult due to the dispersion of biological particles in the air. Peccia and Hernandez (2006) reviewed PCR-based methods that are used in aerosol studies, emphasizing the importance of sample processing, in the DNA extraction steps - cell lysis and purification - as well as in PCR amplification. Longhi et al. (2009) developed a protocol using real-time PCR techniques for pollen identification and obtained promising results even with pollen mixtures. More recently, Kraaijeveld et al. (2015) proposed metabarcoding in combination to NGS as an efficient method to successfully identify pollen taxa using samples from a Hirst-type volumetric device. However, for pollen samples collected by gravimetric traps, no relevant study has been conducted so far.

The aim of this study is to develop protocols for processing environmental samples for DNA extraction and further metabarcoding analysis, and assess the efficacy of these protocols for the taxonomic assignment of airborne pollen, collected by both gravimetric (Tauber trap) and volumetric samplers (Burkard spore trap). Using as a benchmark the classical morphological pollen analysis and building a custom-made reference database, it emerged that DNA metabarcoding is applicable across a complexity of samples, from pure pollen of single species to complex air samples, provided that sample preparation, DNA extraction and amplification protocols are specifically optimized for gravimetric and volumetric sampling.

## 2 Materials and methods

### 2.1 Study area

The sampling area lies in a mountainous region of northern Italy (Trentino), with an elevation between 65 m to 3,764 m above sea level, characterized by diverse phytoclimatic types. The sampling site is in San Michele all’Adige, on the valley floor near the river Adige, 220 m above sea level, in proximity to vineyards and apple orchards, which cover 44% of the area. 8% of the surrounding land is covered by herbaceous weeds like *Parietaria* (*P. diffusa, P. officinalis*), *Artemisia* (*A. absinthium, A. vulgaris, A. verlotorum*) and *Ambrosia* (*Ambrosia artemisiifolia*), while at higher elevations, 48% of the land is forested: 24% mixed (e.g. *Pinus sylvestris, Ostrya carpinifolia, Fraxinus ornus, Quercus pubescens*), 11% coniferous (mainly *Pinus sylvestris, Pinus nigra* and *Picea excelsa*), 8% broad-leaved forests (*Fagus sylvatica, F. ornus, Ostrya carpinifolia, Quercus pubescens,* etc.); the remaining 5% consists of semi-natural areas, transitional woodland, shrubs, and sparsely vegetated areas. (Cristofori et al. 2010).

### 2.2 Sample collection, processing and microscopic analysis

Air samples were collected at the Aerobiological Monitoring Centre of Fondazione Edmund Mach, Trento, Italy by a Hirst-type volumetric trap (VPPS 2000, Lanzoni, Bologna, Italy), which is running at the site since 1988. An additional trap (Sporewatch Electronic Spore & Pollen Sampler, Burkard Manufacturing, Rickmansworth, Hertsfordshire, England) was installed on the same pole at 10 m aboveground. A gravimetric sampling was performed by Tauber traps, two meters far from the volumetric samplers, with the aperture at 0.5 m from the ground. The aperture was equipped with a collar, from which air flows into the trap, covered by a 5 mm mesh net to prevent the deposition of insects and/or other larger particles into the trap. The settling particles were collected in 700 ml of a preservative solution (1:1:1 water, alcohol, glycerol, plus 2 g L^−1^ phenol).

Gravimetric samples were collected over the period 7^th^ of August to 14^th^ of October, 2015 and volumetric samples (from Burkard and Lanzoni traps) were collected over the week 23^rd^ of February to 2^nd^ of March, 2015. Two gravimetric traps and two longitudinal halves (replicates) of Burkard sampling tape were analysed by molecular techniques. Samples of a third gravimetric trap and a Lanzoni weekly tape were analysed by microscopic techniques. A comparison of results from the two different volumetric samplers (Lanzoni and Burkard) was performed over a 5-month period (10^th^ of June to 3^rd^ of November, 2013) in a preliminary study and showed a good correlation of daily pollen data (Spearman’s Rho = 0.91; p<0.001), with no significant differences in terms of pollen load (U Mann-Whitney test). Therefore the pollen data obtained from microscopic analysis of Lanzoni tape can be used as reference for results of Burkard tape molecular analysis.

Pollen pellets for molecular and microscopic analysis were retrieved using the following protocol:

a. Filtration (for Tauber trap samples only): samples were pre-filtered through a 200 μm metal mesh sieve (Retsch, Haan, Germany) to remove large particles (e.g. small insects, plant remains, etc.) and then pollen was collected on a 5 μm mixed cellulose ester filter, 47 mm of diameter (Merck Millipore Ltd, Cork, Ireland), using a vacuum pump filtration system (Sigma-Aldrich, Milan, Italy). When samples were prepared for microscopic analysis, a counting marker (*Lycopodium* spore tablets, Batch 3862, Lund University, Lund, Sweden) was added before filtration. The filters were dried for 2 to 3 h at 65°C, and either immediately processed or stored at −20°C.
b. Pellet preparation: filters or tapes were placed into a glass tube, then 5 ml of acetone were added and vortexed for 2–3 min. By this step the filters were dissolved. After centrifuging (3 min, 2,300 rpm, for each step), the supernatant was discarded and 1 ml of acetic acid was added to the pellet, vortexed and centrifuged as above. After discarding the supernatant, two washing steps were performed to clean the pellet using 2 ml of distilled water, plus one drop of ethanol to reduce the surface tension. The pellet was re-suspended in 1 ml of distilled water, transferred into 2 ml tubes, and centrifuged for 3 min at 13,500 rpm. In case of molecular analysis, the supernatant was discarded and the tubes were stored at −20°C; for microscopic analysis, 0.5 ml of the supernatant was kept, two drops of glycerol were added and the samples were stored at 4°C.

A classic morphological pollen analysis was performed by an optical microscope (Leitz Diaplan, Ernst Leitz Wetzlar, Wetzlar, Germany) under 400x magnification for both tape and Tauber samples. This method gave us a quantitative count of the total number of pollen grains present in each sample during the period of exposure (Table 1). As for Tauber samples, at the end of the pelleting procedure, a small aliquot of the pellet was transferred on a microscope slide and coloured with basic fuchsine. A minimum of 400 pollen grains were analysed at the microscope and the total pollen was then calculated taking into account the number of counted markers (Faegri & Iversen 1989). As for the volumetric sampler (Lanzoni), the tape was mounted on slides, colored with basic fuchsine, and four horizontal continuous sweeps were analysed (14% of the total area), following a national standard (UNI 11108:2004).

**Table 1.** Summary of sample types that were used to develop protocols for processing and analyzing environmental samples by metabarcoding and of their features.

| Sample type | Sample description | Contributing taxa | Complexity factors | Sampling period | No. of pollen grains |
| --- | --- | --- | --- | --- | --- |
| Pure pollen | Samples collected directly from flowers and used for molecular analysis in single species or in mixtures | *Corylus avellana*<br>*Juniperus communis*<br>*Artemisia vulgaris* | none |  | 260,000<br>400,000<br>460,000 |
| Pure pollen on Tape and in Tauber solution | Simulations of volumetric and gravimetric sampling; further used for molecular analysis | *J. communis* | Sampling medium |  | 400,000 |
| E-sample Tape | Volumetric samples collected by Burkard trap and used for molecular analysis in duplicate samples and by Lanzoni trap used for microscopic analysis | Several species (and other bioparticles) | Sampling medium and environmental degradation | 23/02/2015 to 02/03/2015 | 11,500 |
| E-sample Tauber | Gravimetric samples collected by Tauber traps; two traps for molecular analysis and one for microscopic analysis | Several species (and other bioparticles) | Sampling medium and environmental degradation | 07/08/2015 to 14/10/2015 | 79,000 |

### 2.3 DNA extraction experiments

An exploratory experiment was designed to define the DNA extraction conditions yielding the highest amount of DNA from pollen samples. The experimental design was set on the assessment of pollen samples with increasing complexity, from single-species pure pollen to environmental samples (see Table 1 for a summary of all sample types). The starting samples consisted each of 2.5 mg of *Corylus avellana* L., *Juniperus communis* L. and *Artemisia vulgaris L.* pure pollen (directly collected from flowers and stored at 4°C), corresponding to 260,000-460,000 pollen grains, depending on the species, as estimated by a Fuchs-Rosenthal counting chamber (HBG Henneberg-Sander GmbH, Giesen, Germany). The quantity was chosen to reflect the effective amount of pollen grains found in seasonal Tauber traps located within various NATURA 2000 habitat sites, as estimated in a preliminary study (15,000-360,000 pollen grains/Tauber trap, spring season, higher to lower altitudes) using a counting marker.

Tested variables included the pollen disruption method and the DNA extraction kit. A high-energy agitation with beads (Retsch MM200 mixer mill) was applied to help cell lysis by mechanical disruption of cell walls (Retsch); here, glass beads were compared to steel beads. Also, a test was made for the efficiency of cell wall disruption by freezing samples in liquid Nitrogen for 30 s prior to bead beating, as suggested by the DNeasy Plant mini kit protocol (Qiagen GmbH, Hilden, Germany). For each sample, beads were added as follows: 1 steel bead with a diameter of 5 mm (Qiagen) or, alternatively, a mixture of glass beads: 0.3 g of 212-300 μm diameter and 0.2 g of 425-600 μm diameter beads (Sigma-Aldrich). All samples were ground for 1 min at 30 Hz, in two steps.

For the subsequent DNA extraction, two different kits were tested: DNeasy Plant mini kit (Qiagen), which is based on silica gel spin column technology, and Nucleomag kit (Macherey-Nagel, Düren, Germany), which is based on magnetic beads technology. Automated DNA extraction systems were used: Qiacube (Qiagen) and Kingfisher (Thermo Fisher Scientific, Waltham, MA, USA), respectively. The final elution volume was 100 μl.

Taking advantage of the results of the previous steps, DNA was extracted under the so far ‘optimal’ conditions from samples with increasing complexity, simulating pollen grains collected by different means in the field. A fixed quantity of *J. communis* pollen, (2.5 mg), was distributed (i) on a tape (Melinex; DuPont Teijin Films Luxembourg, SA, Luxembourg City, Luxembourg) coated with silicone-based adhesive (Lanzoni) or (ii) in a Tauber trap aqueous solution. Moreover, environmental samples (e-samples), collected by Tauber traps and the Burkard trap in the field and representing complex mixtures of different organisms and particles were analysed. The increasing complexity was first introduced by the sampling medium (tape or aqueous solution) and then by the environmental sampling, because of the co-occurrence, to a varying degree, of biotic particles other than pollen, and of environmental degradation, due to exposure in the field, over different lengths of time (one week for the tape samples and three months for the Tauber ones).

#### 2.3.1 *Statistic analysis*

The DNA yield data were analysed using R software (https://www.r-project.org/). A three-factor ANOVA for all effects and interactions was applied, with the independent variables being the kit, the freezing step and the bead material and the dependent variable the DNA yield. For the extraction under ‘optimal’ conditions, a one-way ANOVA was applied, using the type of sample as independent variable and the DNA yield as dependent variable.

### 2.4 Pollen taxonomic identification

#### 2.4.1 *Local reference database*

The workflow followed for the pollen taxonomic identification of e-samples is summarized in Figure S1. First, a local reference database needed to be constructed. A bioinformatics search was performed in Genbank database (National Center for Biotechnology Information, www.ncbi.nlm.nih.gov/genbank) for a short fragment of the chloroplast trnL intron (trnL c-h barcode) of species known to occur in the study area. In total, 1470 species were searched, belonging to 46 families present in Trentino. Using the R package ‘rentrez’ (https://CRAN.R-project.org/package=;rentrez), trnL sequences were downloaded from the database ‘Nucleotide’ of Genbank using species names as search terms (e.g. ‘*Abies alba* trnL’, Fig. S1.1.a). For each of the sequences, the portion between the primers was retrieved and stored in a database including information on the family, genus, species, sequence identifier number in Genbank (gi) and other metadata. The resulting database was then filtered for wrongly labelled sequences by Genbank. For this, an exhaustive manual validation was done for synonyms of all related species and duplicates were removed. For each family, sequences were then aligned using the ‘Muscle’ algorithm with the R package ‘Bioconductor muscle’ (Edgar 2004). Sequences that could not be aligned due to low quality were removed (Fig. S1.1.b).

Through the previous search, relevant taxa with low or no availability of trnL sequences were identified. For these taxa new sequences were generated. Young leaf samples were collected from different individuals of taxonomically verified plants in San Michele all’Adige (Trentino) and stored at −20°C until DNA extraction. Total DNA was extracted from 100 mg of leaf tissue using the DNeasy Plant mini kit, following the manufacturer’s protocol (Qiagen), and the whole trnL intron was amplified using the primer pair c-A49325 and d-B49863 (Taberlet et al. 2007) (produced by Sigma-Aldrich). Amplicons were Sanger-sequenced in a Genetic Analyzer 3130 (Applied Biosystems, Foster City, CA, USA) and the newly generated sequences were added to already available ones (Fig. S1.1.c), thus forming the final local reference. All sequences stored in the database, together with the related information (i.e. gi, family, genus and species), were exported in a fasta file which was used for the final taxonomic assignment (Fig. S1.1.d).

#### 2.4.2 *PCR Amplification and Sequencing*

Pollen DNA was amplified with c-h trnL primers [5’-CGAAATCGGTAGACGCTACG-3’ and 5’-CCATTGAGTCTCTGCACCTATC-3’] produced by Sigma-Aldrich, and following the Go Taq protocol (Promega, Madison, WI, USA). The DNA amplification was carried out in a final volume of 50 μl using 2 μl of DNA extract as a template. The amplification mixture contained 5 U/μl GoTaq Polymerase, 10x of GoTaq Buffer, 10 mM of dNTPs, 10 μM of each primer, 3 mM of MgCl_2_ and H_2_O. All PCR amplifications were carried out on a Veriti 96 well thermal cycler (Applied Biosystems) with the following program: 2 min at 95°C and 40 cycles of 15 sec at 95°C, 15 sec at 52°C and 30 sec at 72°C. The band sizes and concentrations were checked on a gel electrophoresis using QIAxcel (Qiagen) and analysed by QIAxcel ScreenGel Software.

For samples containing mixtures of species, cloning of PCR products and sequencing of 30 different clones for each PCR product was performed. First, PCR products were run on a 2% agarose gel and bands of the adequate size were excised and purified with the commercial kit QIAquick gel kit (Qiagen). We used the TOPOTA Cloning Kit, Dual Promoter, with pCRII-TOPO Vector and One Shot Mach1 T1 Phage-Resistant Chemically Competent *E. coli* (Invitrogen, Carlsbad, CA, USA) following the manufacturer’s protocol. The DNA amplification of positive colonies was carried out following the Hotmaster Taq protocol (5 Prime GmbH, Hilden, Germany) in a final volume of 20 μl, using 1 μl of cloned DNA as template. The PCR mixture contained 10 μM of each primer M13R and M13F (Sigma-Aldrich), 5 U/μl Hotmaster Taq polymerase, 10x of Hotmaster Buffer, 10 mM of dNTPs and H_2_O. The program was set as follows: 94°C for 2 min, 35 cycles at 94°C for 30 sec, 55°C for 15 sec, 65°C for 2 min and a final elongation step at 65°C for 10 min. b

For all samples, we purified the PCR products using 2 μl of ExoSap-IT (Amersham Pharmacia Biotech, Uppsala, Sweden), 2 μl of H_2_O and 3 μl of the amplified DNA. The purified DNA was then Sanger sequenced using 3.2 μM of the M13 reverse primer for the cloned samples and trnL primers for the rest of the samples and a BrightDye Terminator 2.5X Premix Cycle sequencing mix (Nimagen, Nijmegen, Netherlands). All samples were loaded in a Genetic Analyzer 3130 (Applied Biosystems).

#### 2.4.3 *Taxonomic assignment*

The taxonomic assignment was done with BLAST 2.4.0+ software (downloaded at https://www.ncbi.nlm.nih.gov/news/06-13-2016-blastplus-update/) by comparison of the new unknown sequences to our custom-made local reference database, which was used as a BLASTN database (Fig. S1.2 a). After obtaining the unknown sequences, we used Sequencher version 5.4 sequence analysis software (Gene Codes Corporation, Ann Arbor, MI, USA) in order to assess the quality of the sequences and further process them. All good quality sequences were stored in a fasta file (Fig. S1.2 b). Hence, each of the unknown sequences was aligned to the sequences in the local reference database (Fig. S1.2 c) and information about the alignment was requested (e.g. identity percentage and query coverage) to taxonomically assign the sequences (Fig. S1.2 d).

## 3 Results

### 3.1 Selection of ‘optimal’ protocols

In the DNA extraction test, pure pollen grains from three species, *C. avellana, J. communis* and *A. vulgaris,* were subjected to different extraction protocols to determine the ‘optimal’ DNA extraction conditions leading to the highest DNA yield. The ‘optimal’ conditions were selected by averaging results of the three species, as our aim was to work with complex mixtures of multiple species (environmental samples). Results of the statistical tests and of the obtained DNA yields are shown in Tables 2 and 3, respectively.

**Table 2.** Tested variables for DNA extraction and sample processing protocols. A significant increase in DNA yield (p<0.05) was associated with the selection criterion of the ‘optimal’ extraction conditions; ns = no significant effect.

| Protocols | Tested Variables | Selected variables ($p$-value) | Sample type |
| --- | --- | --- | --- |
| DNA extraction | Freezing Step: with or without | ns | Pure pollen of *Juniperus communis*, *Corylus avellana*, *Artemisia vulgaris* |
| | Bead material: glass or steel | Steel beads ($p < 0.001$) | |
| | Extraction kit: DNeasy Plant mini or Nucleomag | Nucleomag kit ($p < 0.05$) | |
| Sample processing | Extraction kit: DNeasy Plant mini or Nucleomag + Washing steps: 2 or 4 | Nucleomag kit + 4 washing steps, Pure pollen in Tauber solution ($p < 0.001$) | Pure pollen of *J. communis* on Tape and in Tauber solution |

**Table 3.** DNA yields under optimal conditions (i.e. steel beads, Nucleomag kit, four washing steps) for pure pollen of three different taxa, for *J. communis* on tape and in Tauber solution, and for environmental samples collected in Tauber solution and on tape.

| Species | Sample type | DNA yield (ng/ul)<br>(± sd) |
| --- | --- | --- |
| *Artemisia vulgaris* | Pure pollen | 2.66 ± 1.84 |
| *Corylus avellana* | Pure pollen | 0.31 ± 0.14 |
| *Juniperus communis* | Pure pollen | 18.56 ± 11.01 |
| *J. communis* | Pure pollen in Tauber solution | 8.8 ± 4.39 |
| *J. communis* | Pure pollen on tape | 3.1 ± 3.16 |
| | E-sample Tauber | 1.36 ± 0.21 |
| | E-sample tape | 0.2 ± 0.02 |

The DNA yield, analysed by three-factor ANOVA, showed statistically significant differences for the DNA extraction kit (p<0.05) and the bead material (p<0.001). In particular, the highest DNA yield was obtained with the ‘Nucleomag’ extraction kit and a disruption step performed with steel beads; adding a freezing step by liquid nitrogen did not influence the DNA yield. Finally, the ANOVA test showed no significant interactions between the variables examined.

When applying the same extraction protocol to pure pollen of *J. communis* after simulating sample collection on tape (as in volumetric sampling) or in Tauber solution (as in gravimetric sampling), the DNA yield decreased significantly in both cases (p<0.05). In order to improve the DNA yield, a second experiment was performed to examine if the result was influenced by chemicals used in the sample processing steps (i.e. negative interactions with chemicals of the extraction kit and PCR inhibition). Using *J. communis* pollen, we examined whether for samples extracted by Qiagen kit, PCR was less influenced by the sample processing or if additional washing steps would increase DNA yield (Fig. 1). A three-way ANOVA was performed for factor interactions. In the case of Tauber solution samples, there was a significant increase in DNA yield when the washing steps were doubled (p<0.001); no analogous significant difference was found for the tape samples.

**Fig. 1.**
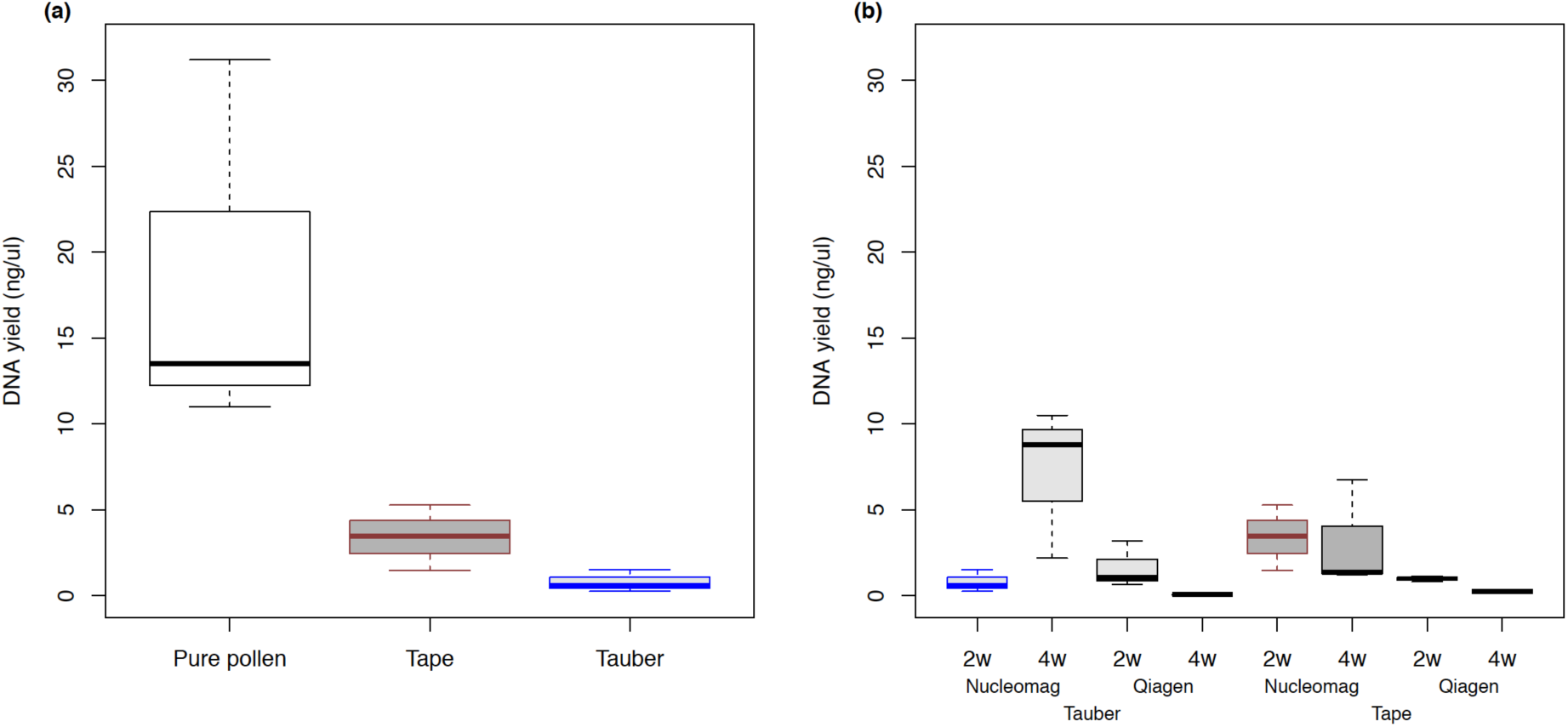
DNA yield for the different types of samples of *J. communis.* The boxplots give the 25^th^ and 75^th^ percentile (bottom and top of the box, respectively) and the median (black, brown and blue thick line). The upper and lower whiskers correspond to the maximum and minimum DNA yield values, respectively. Boxplot (a) provides DNA yields of the extraction for pure pollen (white box), pollen from tape (grey box) and Tauber solution (light grey box) after two washing steps, Nucleomag kit and steel beads. Boxplot (b) provides DNA yield of pollen from tape (grey box) and Tauber (light grey box) samples after two (2w) and four (4w) washing steps for two extraction kits (Nucleomag and Qiagen) and steel beads.

As a result, the ‘optimal’ conditions were defined as those including disruption of the pollen cell wall with steel beads, DNA extraction with the Nucleomag kit and four washing steps during the sample processing.

### 3.2 Taxonomic identification

After a bioinformatics search for the trnL c-h barcode, a total of 1188 sequences were retrieved, corresponding to 403 species of 198 genera and 46 families; 44 sequences, corresponding to 26 species, were newly generated (Table S1).

For single species pollen, the extraction test resulted in a correct taxonomic assignment for all examined samples. All *J. communis* tape or Tauber samples were successfully sequenced and taxonomically assigned. Finally, for the environmental samples, 124 positive clones were recovered from the tape and 117 clones from the Tauber solution. 31 and 28 amplicons, respectively, were selected to be sequenced. The sequences were identified using BLASTN 2.4.0+ against the custom-made local reference database. All sequences were identified with a 100% identity match on the whole sequence length (100% query coverage), except for *Urtica dioica* whose taxonomic identity was 99.2%, with a coverage of 99%. The taxonomic assignment resulted in 7 different taxa for the tape and 3 different taxa for the Tauber solution. When compared to microscopic results (Table 4), 75% and 37% of the identified taxa by microscope were revealed by the molecular method for the volumetric and the gravimetric sampling, respectively. *Cedrus* was dominant in the sequencing results of the Tauber environmental sample.

**Table 4.**
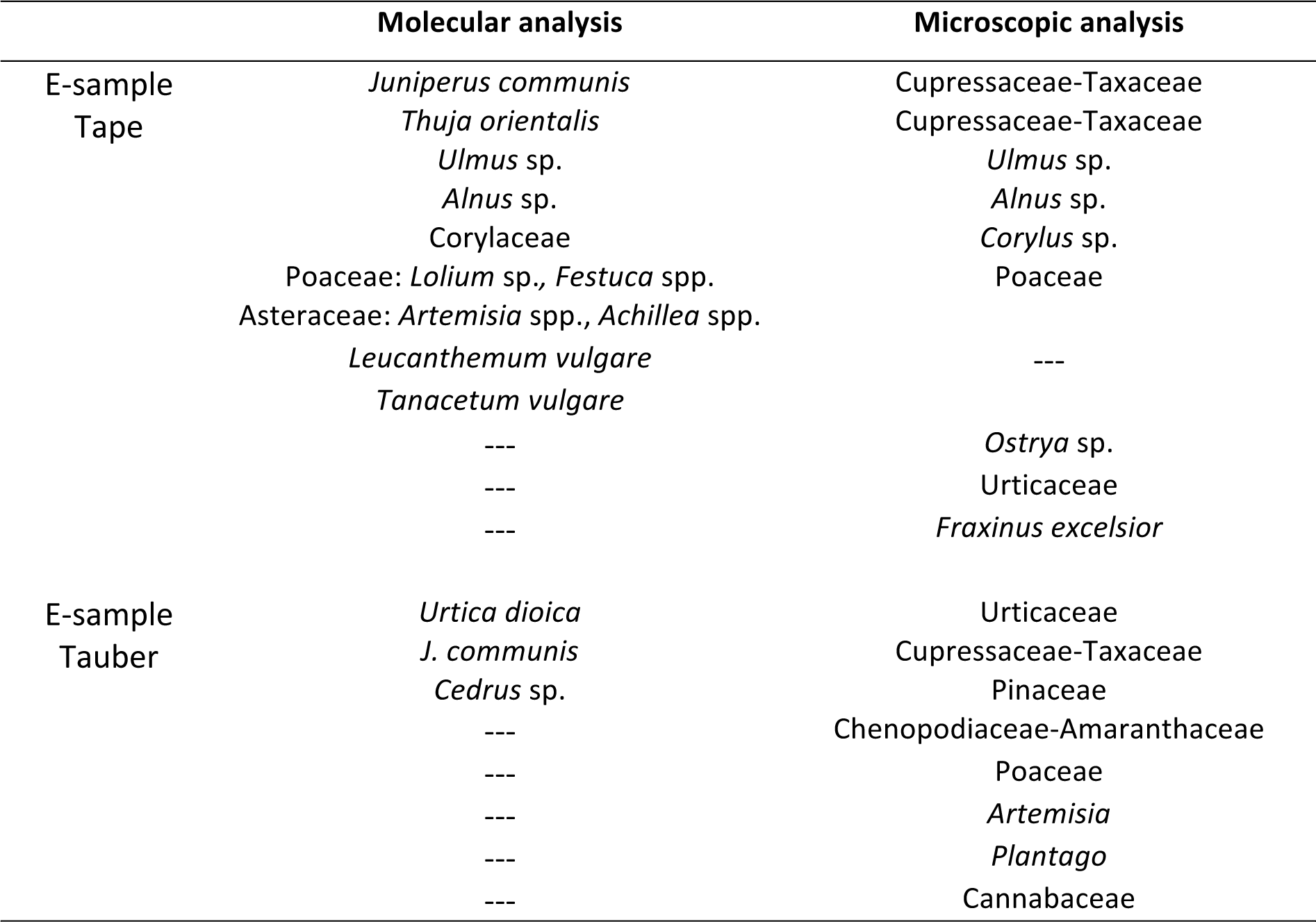
Taxa identified by molecular and microscopic analyses (reference taxa) for the environmental samples (e-samples) collected on the aerobiological tape (replicate halves of tape) and the Tauber trap (replicate traps).

The molecular method provided on average a higher taxonomic resolution. For example, pollen identified microscopically as Cupressaceae/Taxaceae was assigned to *J. communis* and *Thuja orientalis* by molecular analysis. For big families, such as Poaceae and Asteraceae, the molecular method gave 2 and 5 assigned taxa, respectively. On the contrary, within the Corylaceae family, the trnL could not discriminate among *Ostrya carpinifolia, Carpinus betulus* and *C. avellana,* whereas the microscopic analysis could assign pollen to the species level. For all taxa, the molecular method was able to assign sequences at least to the family level; this was not the case for the microscopic method which could not distinguish Chenopodiaceae from Amaranthaceae and Cupressaceae from Taxaceae.

## 4 Discussion

In this study, a DNA extraction protocol for environmental samples of pollen was determined in order to maximize the DNA yield. The method worked properly for all pollen types, although the DNA yield was different across taxa; this is probably related to the cell wall structure (Kraaijeveld et al. 2015). For instance, we recorded a higher DNA yield for *J. communis.* This species is characterized by a cell wall with a very thin layer of exine, which allows an easier disruption of the cell and subsequent release of DNA. Apart from the lysis, all the remaining steps can be easily automated (automatic DNA extraction instrument), making the procedure fast and cost-effective.

Comparing the two pollen collecting methods and using the same quantity of *J. communis* pollen, we found that the total amount of DNA recovered was significantly higher in the Tauber solution, when adding the extra washing steps, which improved the quality and quantity of DNA extracted from the Tauber solution. This suggests that residues in the different steps of the procedure to obtain the pellet affected negatively the DNA extraction, probably due to interactions between the medium of the gravimetric trap and kit chemicals.

The DNA extraction conditions that we selected were finally tested on environmental samples. They resulted into lower DNA yield compared to that of pure pollen grains and pure pollen grains from tape or Tauber samples. This might be due to at least three reasons: (i) the concentration of pollen grains in the environmental samples was lower, (ii) environmental degradation is possible, and (iii) environmental samples contain several other biological particles (e.g. fungi, bacteria, etc.) that may negatively affect the DNA extraction steps. For the sample preparation, we tried to minimize material loss; the type of filter that we selected gets easily dissolved in acetone, the tape was also eluted with acetone, and all the material remained in the tube until the step of extraction.

Before any molecular-based taxonomic assignment, it is fundamental to have a comprehensive reference database. To this aim, we used a fragment of the trnL (c-h), already adopted for pollen metabarcoding (Kraaijeveld et al. 2015). We, thus, carefully built a local reference database integrating available sequences with newly acquired ones, so as to have sequences from the most representative plant species of the region.

Sanger-sequencing on pure pollen grains resulted in a correct taxonomic assignment on all samples of this type. For environmental samples, we adopted a semi-metabarcoding approach, by using the traditional molecular-based approach of cloning and sequencing of several amplicons of the PCR product. This allows to partially overcome the limit of Sanger sequencing, which can only sequence single species, individually. We used for comparison the results of classical morphological identification of pollen grains from replicate environmental samples. In the case of the aerobiological tape samples, the c-h barcode identified the majority of the taxa that were determined by morphological analysis, whereas for Tauber solution samples it was less efficient. However, in both cases, the DNA barcode provided higher taxonomic resolution, with the exception of *C. avellana.* For instance, the morphological analysis assigned pollen to Cupressaceae/Taxaceae and Poaceae families while the molecular analysis assigned them to *J. communis* and *T. orientalis,* and to *Lolium/Festuca,* respectively. This demonstrates that molecular analysis is particularly important for families where pollen morphology is conserved across different genera and species, such as Cupressaceae/Taxaceae. Our results, even though based on Sanger sequencing, confirmed a resolution at the genus level for grasses as showed by Kraaijeveld et al. (2015).

Indeed, the most effective approach to take advantage of the efficiency of DNA metabarcoding is determining a large number of different sequences, which is the main aim of our study, by means of NGS. The successful outcome of any NGS approach is highly dependent on DNA quality and quantity, i.e. sufficient amounts of high-quality nucleic acids, which will be representative of the pollen cells in the sample, for subsequent library preparation (Bell et al. 2016). This is the reason why we designed experiments for developing protocols to obtain high quality pollen DNA from environmental samples. We admittedly used a suboptimal methodology for taxonomic assignment, i.e. Sanger-sequencing cloned PCR products from each environmental sample. Nonetheless, the final outcome is encouraging: good DNA quality can be efficiently retrieved from a complex mixture of pollen grains and other particles. Moreover, as expected, the molecular analysis allowed higher taxonomic resolution compared to microscopic analysis.

The most critical issue is the low efficiency in the identification of pollen from gravimetric samples: only 37% of the morphologically identified taxa were detected. The volumetric sampling has advantages over the gravimetric one regarding the DNA quality as the tape is exposed in the field only for a short period, one week, whereas the Tauber traps remain for longer, three months in our experiments. Moreover, the aqueous medium of the Tauber trap can lead to increased DNA degradation compared to the tape of the volumetric sampler, which is an inert medium where particles attach. The medium of the gravimetric sampler is expected to be more prone to oxidative reactions and microorganism proliferation resulting in higher rates of DNA degradation and accumulation of molecules other than those of pollen. Secondary metabolites, produced by degradation of biological particles, may interact negatively with chemicals used in DNA extraction and amplification. Finally, since the Tauber traps are closer to the ground level it cannot be excluded that more particulate matter is collected in the medium. This is another factor potentially affecting DNA amplification. The scarce representation of taxa found by molecular analysis in the Tauber solution may also depend on the taxa collected. It is known (Parducci et al. 2013) that different pollen grains may carry a different number of chloroplasts; in particular, most of the conifers are reported to be rich in cpDNA, because of its exceptional paternal inheritance (Bennett & Parducci 2006). This might partially explain the predominance of *Cedrus* in our Tauber trap sample sequencing results.

Several lines of evidence (Bell et al. 2016) suggest that a NGS-based metabarcoding approach could significantly increase the information from environmental samples in terms of number of identified taxa. Also, for a better taxonomic assignment in families, such as Poaceae and Asteraceae, the use of an additional DNA barcode, preferably a nuclear one like ITS may be very efficient (De Barba et al. 2014). Parducci et al. (2013) argued that metabarcoding and microscope-based methods can be considered as complementary in revealing different taxa. Our results showed that metabarcoding is a very useful method for taxonomic assignment of pollen grains retrieved from environmental samples, but that they cannot entirely substitute the classical microscopic techniques, particularly in the case of gravimetric samples. This is where more research is needed for attaining higher resolution of the molecular method.

## Acknowledgements

This project is supported by FIRS>T (FEM international Research School). We thank Maria Cristina Viola for microscopic analyses and for helping with the taxonomic identification of plants. We also thank Conservation Genetics group of FEM Research and Innovation Centre (Matteo Girardi, Barbara Crestanello, Andrea Gandolfi, Alice Fietta) and the Zooplant Lab group of Milano Bicocca University (Maurizio Casiraghi, Antonella Bruno and Anna Sandionigi) for support in molecular techniques, Camilla Capelli, Athanasios Charalampopoulos and Duccio Rocchini for their precious suggestions.

## FIGURE LEGEND

**Fig. S1.**
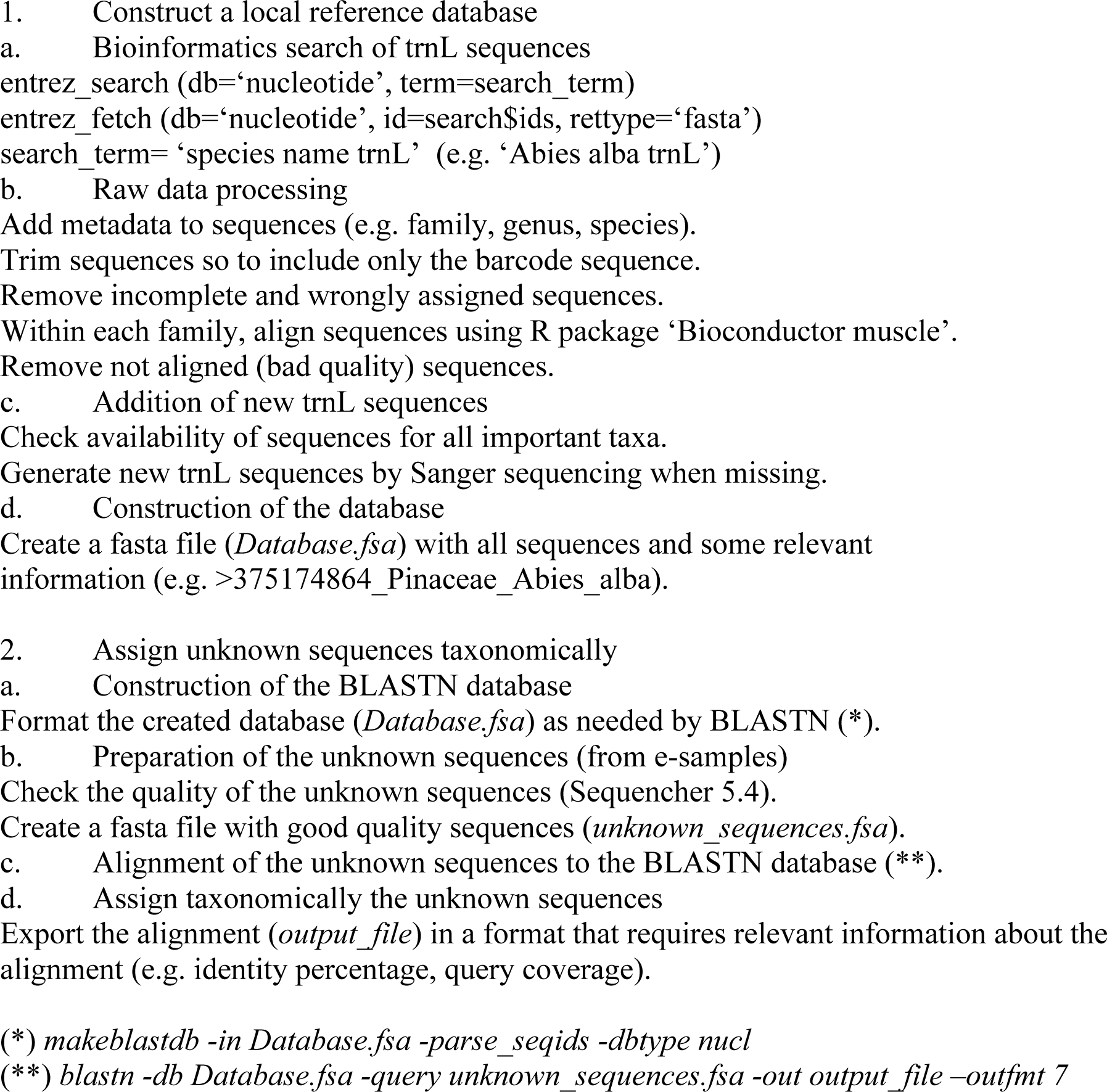
Workflow followed for pollen taxonomic identification of environmental samples, including the construction of a local reference database and the taxonomic assignment of unknown sequences of pollen taxa.

**Table S1.** List of species for which new trnL sequences were generated with Sanger sequencing and their accession number in Genbank. Where different accession numbers are present for the same species, they refer to different individuals (e.g. *Amaranthus retroflexus,* trnL sequence from 3 individuals).

| Species | Family | Accession number (Genbank) |
| --- | --- | --- |
| *Amaranthus retroflexus* | Amaranthaceae | KY313859, KY313860, KY313861 |
| *Hedera helix* | Araliaceae | KY313862, KY313863, KY313864 |
| *Artemisia vulgaris* | Asteraceae | KY313902 |
| *Ambrosia artemisiifolia* | Asteraceae | KY313896 |
| *Betula pendula* | Betulaceae | KY313865, KY313866 |
| *Buxus sempervirens* | Buxaceae | KY313867, KY313868, KY313869 |
| *Humulus lupulus* | Cannabaceae | KY313870 |
| *Sambucus nigra* | Caprifoliaceae | KY313871 |
| *Ostrya carpinifolia* | Corylaceae | KY313872, KY313873 |
| *Corylus avellana* | Corylaceae | KY313894, KY313895 |
| *Thuja orientalis* | Cupressaceae | KY313897 |
| *Juniperus communis* | Cupressaceae | KY313898 |
| *Cupressus sempervirens* | Cupressaceae | KY313874 |
| *Cupressus arizonica* | Cupressaceae | KY313899 |
| *Ginkgo biloba* | Ginkgoaceae | KY313875 |
| *Juglans regia* | Juglandaceae | KY313876 |
| *Laurus nobilis* | Lauraceae | KY313877, KY313878 |
| *Olea europaea* | Oleaceae | KY313879, KY313880 |
| *Fraxinus ornus* | Oleaceae | KY313881, KY313882 |
| *Trachycarpus fortunei* | Palmaceae | KY313883, KY313884 |
| *Larix decidua* | Pinaceae | KY313885, KY313886 |
| *Cedrus* sp. | Pinaceae | KY313887 |
| *Platanus* sp. | Platanaceae | KY313900 |
| *Aesculus hippocastanum* | Sapindaceae | KY313888, KY313889 |
| *Ulmus* sp. | Ulmaceae | KY313901 |
| *Parietaria* sp. | Urticaceae | KY313890, KY313891, KY313892,<br>KY313893 |

## References

Bell, K. L., Vere, N. De, Keller, A., Richardson, R. T., Gous, A., Burgess, K. S., & Brosi, B. J. (2016). Pollen DNA barcoding : current applications and future. Genome 59(9), 1–12. doi: 10.1139/gen-2015-0200

Bennett, K. D., & Parducci, L. (2006). DNA from pollen : principles and potential. The Holocene, 16(8), 1031–1034. doi: 10.1177/0959683606069383

Cristofori, A., Cristofolini, F., & Gottardini, E. (2010). Twenty years of aerobiological monitoring in Trentino (Italy): Assessment and evaluation of airborne pollen variability. Aerobiologia, 26(3), 253–261. doi:10.1007/s10453-010-9161-3

Damialis, A., Halley, J. M., Gioulekas, D., & Vokou, D. (2007). Long-term trends in atmospheric pollen levels in the city of Thessaloniki, Greece. Atmospheric Environment, 41(33), 7011–7021. doi:10.1016/j.atmosenv.2007.05.009

De Barba, M., Miquel, C., Boyer, F., Mercier, C., Rioux, D., Coissac, E., & Taberlet, P. (2014). DNA metabarcoding multiplexing and validation of data accuracy for diet assessment: Application to omnivorous diet. Molecular Ecology Resources, 14(2), 306–323. doi:10.1111/1755-0998.12188

Dell’Anna, R., Cristofori, A., Gottardini, E., Monti, F. (2010). A critical presentation of innovative techniques for automated pollen identification in aerobiological monitoring networks. In B.J. Kaiser Pollen: Structure, types and effects, (pp. 273–288). New York: Nova science

Edgar, R.C. (2004) MUSCLE: multiple sequence alignment with high accuracy and high throughput. Nucleic Acids Research 32(5), 1792–1797. doi:10.1093/nar/gkh340

Faegri, K., & Iversen, J. (1989). Textbook of pollen analysis. London: Wiley

Kraaijeveld, K., de Weger, L. A., Ventayol García, M., Buermans, H., Frank, J., Hiemstra, P. S., & den Dunnen, J. T. (2015). Efficient and sensitive identification and quantification of airborne pollen using next-generation DNA sequencing. Molecular Ecology Resources, 15(1), 8–16. doi:10.1111/1755-0998.12288

Levetin, E. (2004). Methods for aeroallergen sampling. Current allergy and asthma reports, 4(5), 376–383. doi:10.1007/s11882-004-0088-z

Longhi, S., Cristofori, A., Gatto, P., Cristofolini, F., Grando, M. S., & Gottardini, E. (2009). Biomolecular identification of allergenic pollen: A new perspective for aerobiological monitoring? Annals of Allergy, Asthma and Immunology, 103(6), 508–514.doi:10.1016/S1081-1206(10)60268-2

Oteros, J., Pulsch, G., Weichenmeier, I., Heimann, U., Möller, R., Röseler, S., et al. (2015). Automatic and Online Pollen Monitoring International Archives of Allergy and Immunology, 167(3), 158–166. doi:10.1159/000436968

Parducci, L., Matetovici, I., Fontana, S. L., Bennett, K. D., Suyama, Y., Haile, J., et al. (2013). Molecular- and pollen-based vegetation analysis in lake sediments from central Scandinavia. Molecular Ecology, 22(13), 3511–3524. doi:10.1111/mec.12298

Peccia, J., & Hernandez, M. (2006). Incorporating polymerase chain reaction-based identification, population characterization, and quantification of microorganisms into aerosol science: A review. Atmospheric Environment, 40(21), 3941–3961. doi:10.1016/j.atmosenv.2006.02.029

Taberlet, P., Coissac, E., Pompanon, F., Gielly, L., Miquel, C., Valentini, A., Vermat, T., Corthier, G., Brochmann, C., Willerslev, E. (2007). Power and limitations of the chloroplast trnL (UAA) intron for plant DNA barcoding. Nucleic acids research, 35(3), e14–e14.

